# Plant nutritional and metabolic responses to drought and elevated CO_2_ revealed by machine learning-enabled non-targeted metabolomics

**DOI:** 10.1101/2025.01.02.631073

**Authors:** Hsin-Fang Chang, Theresa Caso-McHugh, David L. Des Marais

## Abstract

Projected atmospheric CO_2_ rise, coupled with intensification of drought in many regions, impacts the physiology of C_3_ plants beyond photosynthesis and carbon metabolism. The interaction between CO_2_ and drought affects the concentrations of many nutrients in crops, often resulting in excessive agrochemical use and less nutritious food production. To address these challenges, we investigated nutrient dynamics in *Brachypodium distachyon*, a model crop plant, under ambient and elevated CO_2_, factorially combined with well-watered or drought treatments. Integrative analyses of plant physiology, transcriptomics, and machine learning-enabled non-targeted metabolomics revealed that plant elemental composition and metabolomic responses to elevated CO_2_ strongly depend on water availability and differ between shoots and roots. Elevated CO_2_ and drought significantly impaired nitrogen status, with root nitrate uptake being more negatively affected than ammonium uptake. However, elevated CO_2_ increased iron partitioning in shoots under drought, potentially driven by enhanced carbon availability facilitating chelator synthesis for iron translocation. The high accumulation of sphingolipids in roots under combined stresses suggests a protective role against ionome imbalances. These findings highlight how climate stressors interact to shape plant nutrient dynamics, providing insights that can guide agricultural practices and breeding strategies to optimize nutrient management and foster sustainable agriculture under changing climate.

## Introduction

Atmospheric CO_2_ concentrations have been rising since the onset of the Industrial Revolution, a trend projected to continue alongside increasing incidences of low water availability (Lee et al., 2023). Drought stress already represents a major constraint on crop yield (Nemani et al., 2003; Cattivelli et al., 2008; Farooq et al., 2009) and nutritional quality (Peterson et al., 1992; Champolivier and Merrien, 1996; Singh et al., 2008; Vandegeer et al., 2012). Over the past four decades, global droughts have caused an estimated 1820 million Mg of cereal crop losses(Lesk et al., 2016), with future impacts expected to intensify under rising CO_2_ levels (Li et al., 2009; Gray et al., 2016). Understanding the mechanisms of interactions between crops and climate stressors may facilitate new breeding and management strategies to mitigate impacts of climate change on agriculture.

When faced with water limitation, plants need to maintain turgor, stabilize membrane permeability, and maintain water status through osmotic adjustments (Wang et al., 2013). Stomatal closure, a primary response to drought, serves as the starting point for transpiration reduction. Reduced transpiration reduces mass flow, thereby affecting nutrient translocation from roots to shoots and, thus, root uptake capacities (Hu et al., 2007; Farooq et al., 2009). Decreases in transpiration resulting from elevated CO_2_ levels may further impact transport. Debates over whether future CO_2_ levels will have an adverse effect on the nutrient statuses of crops are ongoing. Recent meta-analyses of free air CO_2_ enrichment experiments have linked elevated CO_2_ to reductions in nutrients, including iron and zinc, in C_3_ crops (Myers et al., 2014), with carbon dilution (i.e., biomass fertilization effect) or reduction in mass flow suggested as possible explanations (Taub and Wang, 2008; Mcgrath and Lobell, 2013).

Environmentally dependent changes in plant nutrient assimilation and subsequent altered elemental composition and stoichiometry of their tissues should be observable through metabolomic analysis. Changes in plant elemental composition and stoichiometry are linked with shifts in metabolism, responding to both abiotic (Rivas-Ubach et al., 2012) and biotic factors(Wu et al., 2013). In plants exposed to drought, proline, sugars(Hoekstra et al., 2001), and the osmoprotectants choline and glycine betaine accumulated in shoots, whereas disaccharides, amino acids, and elements involved in growth (potassium) accumulated in roots (Gargallo-Garriga et al., 2015). However, we know little of the relationships between the overall use of nutrients by plants, and shifts in metabolites under the combined stresses of drought and elevated CO_2_—a critical knowledge gap in light of anticipated climate change.

In this study, we grew *Brachypodium distachyon* (hereafter Brachypodium) under one of two watering regimes, and exposed them to either ambient CO_2_ (400 ppm) or elevated CO_2_ (800 ppm) environments. We chose Brachypodium, a model C_3_ grass species, because it shares close phylogenetic relationships and high genomic synteny with temperate cereal crops(Catalan et al., 2016). As a genetically diverse undomesticated species with a wide geographical distribution, Brachypodium is well-suited for research into stress response mechanisms. An integrated -omics analysis—encompassing the ionome, transcriptome, and non-targeted metabolome—was used to test the hypothesis that elevated CO_2_ differentially influences the effects of drought on elemental composition and metabolomics in shoots and roots.

## Results

### Effects of drought and elevated CO_2_ levels on leaf gas-exchange parameters

Drought suppressed the net photosynthesis rate (*A*_n_) of Brachypodium and this decrease was greater under elevated CO_2_ (-75%) than ambient CO_2_ (-45%) treatments (Figure 1a). Stomatal conductance (*g*_s_) was lower under drought, which was further accentuated by elevated CO_2_ (Figure 1b). Under drought alone, transpiration was reduced by 62.8%, while under the combined effects of drought and elevated CO_2_, it decreased by 93.8% (Figure 1c). Drought increased intrinsic water use efficiency (iWUE; ratio of *A_n_* to *g*_s_; Figure 1e) and δ^13^C, (Figure 1f) which is an integrated measure of WUE in C_3_ plants(Farquhar et al., 1989); this increase tended to be greater under elevated CO_2_ than ambient CO_2_ conditions. The ratio of intercellular to ambient CO_2_ concentration (C_i_/C_a_) decreased under drought and this pattern was maintained under elevated CO_2_ levels (Figure 1d).

**Figure 1.**
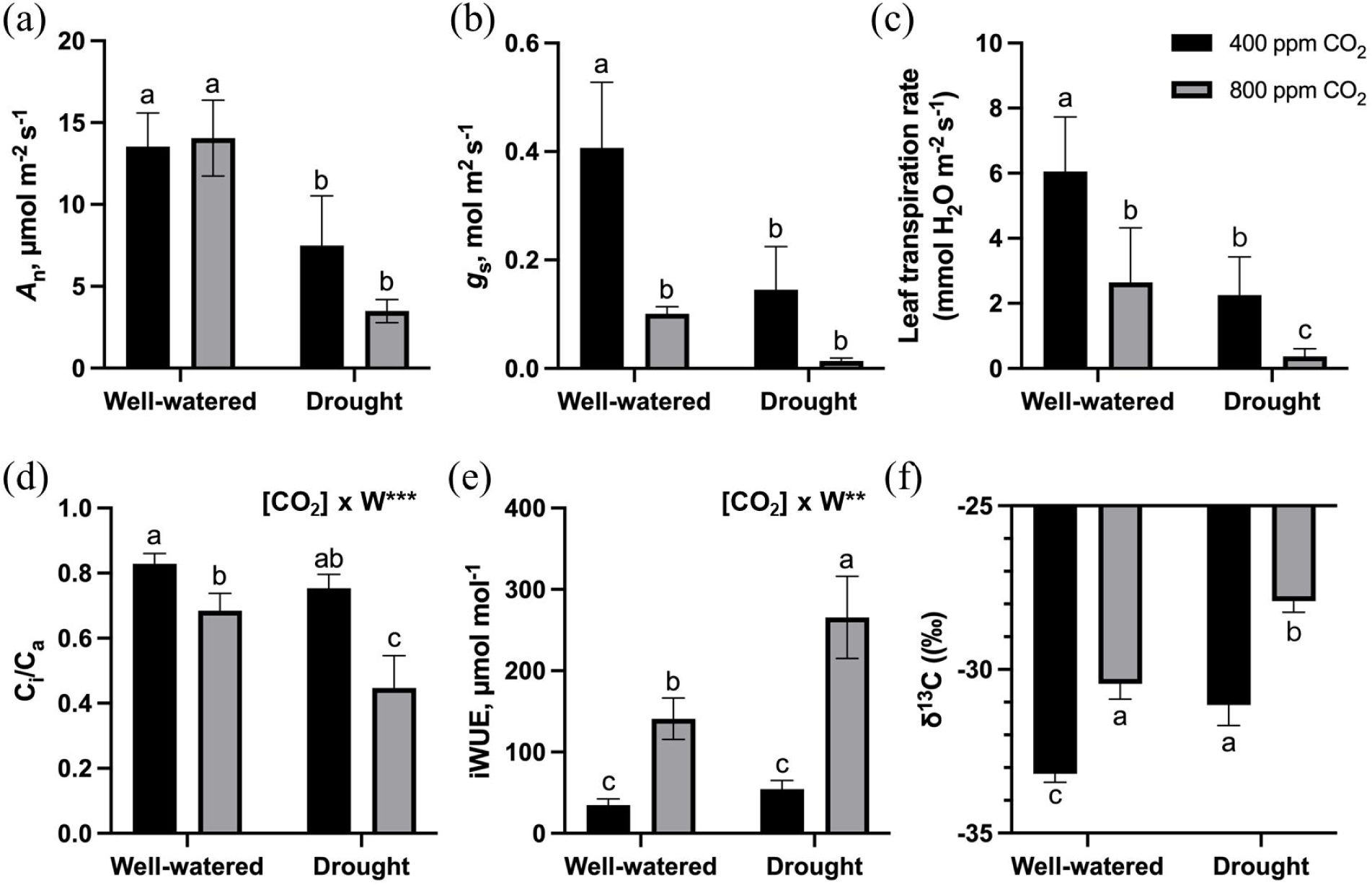
Leaf gas-exchange parameters. (a) Net photosynthesis rate (*A*_n_), (b) stomatal conductance (*g*_s_), (c) transpiration rate, (d) instantaneous photosynthetic water use efficiency (iWUE), (e) intercellular (C_i_) to ambient (C_a_) CO_2_ concentration ratio, and (f) WUE as determined by δ^13^C. Lower (more negative) values of δ^13^C indicate lower WUE. Plants were grown under 400 ppm CO_2_ (ambient CO_2_) or 800 ppm CO_2_ (elevated CO_2_) and two water (W) regimes (well-watered and drought). Each bar represents mean ± SD (n = 3). Different lowercase letters indicate *P* < 0.05 by the least significant difference test. W × CO_2_ effect; ns, *P* > 0.05, not significant; \**P* < 0.05; \*\**P* < 0.01; \*\*\**P* < 0.001; \*\*\*\**P* < 0.0001.

Under well-watered conditions, the plants grown under elevated CO_2_ had a lower maximum carboxylation rate of Rubisco (*V*_c,max_) as compared to control CO_2_ and exhibited no depression of maximum electron transport rate, leading to ribulose-1,5 *bis*phosphate (RubP) regeneration (*J*_max_) (Figure S1). In drought-treated plants, there was a trend towards an acclimatory response to elevated CO_2_ as demonstrated by lower (-73%) *V*_c,max_ and lower (-87%) *J*_max_ compared to ambient CO_2_-grown plants. As a result, the ratio of *J_max_* to *V*_c,max_ was greater under drought and elevated CO_2_ conditions. Respiration (R_d_) rate, as approximated from A/C_i_ curves (Figure S2), was stimulated by elevated CO_2_ (+82%) when plants were well-watered, but decreased by 55% under drought (Figure S1). Neither elevated CO_2_ nor soil drying had a significant effect on *Fo*, *Fv/Fm*, *PhiPSII*, *qP*, *NPQ* and *qL* (all *p* > 0.05; Figure S3).

### Water stress and elevated CO_2_ levels impact plant growth and the uptake, accumulation, and partitioning of C and nutrients

Drought decreased the shoot and root biomass at elevated CO_2_, while only increasing root biomass at ambient CO_2_ (Figure S4). Total biomass of plants was increased under elevated CO_2_ when plants were well-watered, but decreased at high CO_2_ under drought conditions, indicating that drought negated the stimulation of plant growth by elevated CO_2_ level (Figure S4). Regardless of the watering regime, elevated CO_2_ increased C concentrations in the shoot, but had no effect on root C (Figure 2). Compared to control watering and to ambient CO_2_, both drought and elevated CO_2_ decreased shoot N concentration by nearly half. Conversely, both elevated CO_2_ and drought more than doubled the ratio of C to N, in both shoots and in roots (Figure 2).

**Figure 2.**
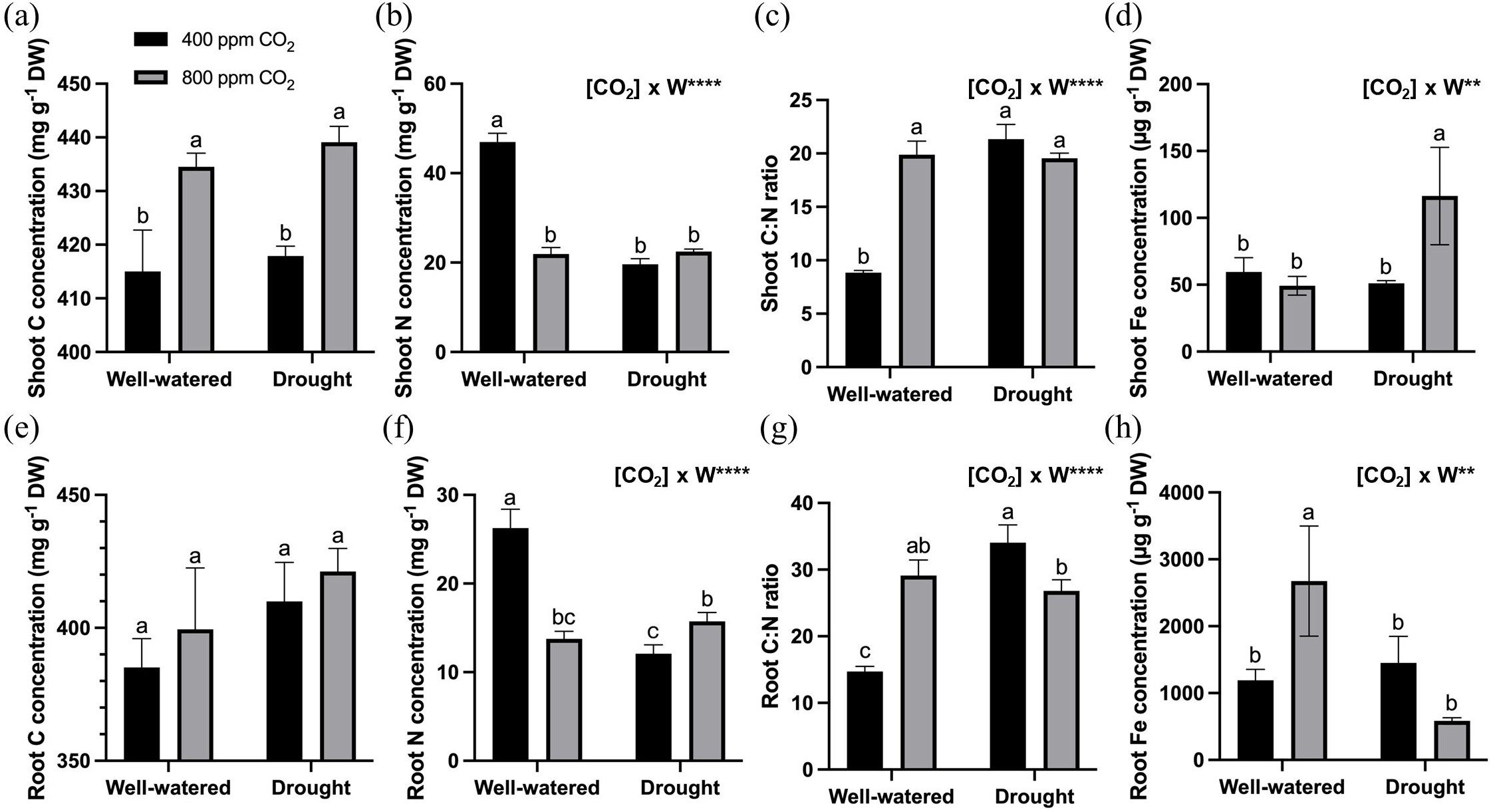
Plant C and nutrient concentration. (a), (e) C concentrations, (b), (f) N concentrations, (c), (g) C:N ratio, and (d), (h) Fe concentrations in either shoots or roots, respectively, of Brachypodium. Plants grown under 400 ppm CO_2_ (ambient CO_2_) or 800 ppm CO_2_ (elevated CO_2_) and two water (W) regimes (well-watered and drought). Each bar represents mean ± SD (n = 3). Different lowercase letters indicate *P* < 0.05 by the least significant difference test. W × CO_2_ effect; ns, *P* > 0.05, not significant; \**P* < 0.05; \*\**P* < 0.01; \*\*\**P* < 0.001; \*\*\*\**P* < 0.0001.

Drought slightly, but not significantly, decreased Ca concentrations in shoots, while shoot Ca concentration was strongly reduced by elevated CO_2_ under well-watered conditions (Figure S6a). Patterns of root Ca are more complex, with well-watered control CO_2_ roots having the lowest concentration and droughted high CO_2_ roots having the highest (Figure S6e). Interestingly, these are the conditions with the highest and lowest stomatal conductance, respectively (Figure 1b). We observe the reverse pattern for root K concentrations, in which concentrations are the lowest under drought plus elevated CO_2_ (Figure S6g). Both drought and elevated CO_2_ decrease shoot Ca concentrations to a similar degree. Shoot Mg, Mn, and Cu were not affected by water regimes and CO_2_ levels. However, drought decreased Mg and Mn in roots, while Cu remained unchanged. In shoots, Fe, Zn, and Na concentrations were higher under drought and elevated CO_2_, resulting in a significant CO_2_ × Water interaction. Drought decreased root Fe concentration, while increasing root Zn and having no effect on Na (Figure 2; S6).

Total uptake of all macro- (Ca, Mg, K, Na, P) and micronutrients (Fe, Mn, Cu, Zn), except for N, increased under elevated CO_2_ (Figure S5; S8). Conversely, combined drought and elevated CO_2_ stresses led to a decrease in the uptake of all these nutrients (Figure S5; S7). There were large differences in partitioning of nutrients between the root and shoot. Under elevated CO_2_, the allocation of Ca, K, and Zn to the shoot remained unchanged. However, more N remained in the root under drought combined with elevated CO_2_, while this combination suppressed N partitioning to the shoot (Figure S5). Under these conditions, greater partitioning of Mg, K, Na, Fe, and Mn to the shoots was observed (Figure S8).

### Transcriptome responses in plants subjected to the combination of drought and elevated CO_2_ show unique patterns

Compared to the control group, the drought and elevated CO_2_ treatments resulted in 2201 and 1004 DEGs in shoots, and 5156 and 281 DEGs in roots, respectively (Figure S9b). Under the combined stresses, 737 DEGs in shoots and 781 DEGs in roots were uniquely affected by the interaction of drought and elevated CO_2_.

Hierarchical clustering of DEGs revealed six clusters in each tissue, plotted here as the average abundance of genes in each cluster (Figure S10). In shoots, clusters 1 and 2 largely characterize positive or negative response to drought stress, and show the apparently ameliorating effect of elevated CO_2_ on response to drought. Conversely, the expression of genes in clusters 3 and 5 exhibit an enhanced response of drought under elevated CO_2_. In roots, clusters 5 and 3 contained genes that were upregulated and downregulated, respectively, in response to drought at both ambient and elevated CO_2_ levels (Figure S10). Details of the GO and KEGG analyses are provided in section S2.2 of the Supporting Information.

### Expression of transport-associated genes and known ionome genes in relation to ionomic changes under water deficit and elevated CO_2_ levels

There were 23 and 27 DEGs with GO-annotated transport functions or identified as known ionome genes (KIGs(Whitt et al., 2020)), in shoots and roots, respectively (Figure 3). Among them, 42% were related to the transport of a broad group of nutrients comprising Fe, Mn, Cu, and Zn, and a third to N transport, while the remaining were linked to the transport of Ca, Mg, K, and Na. Although these proportions reflect the number of genes tagged for each element, there are contrasting trends when focusing on transporter genes and KIGs related to each nutrient.

**Figure 3.**
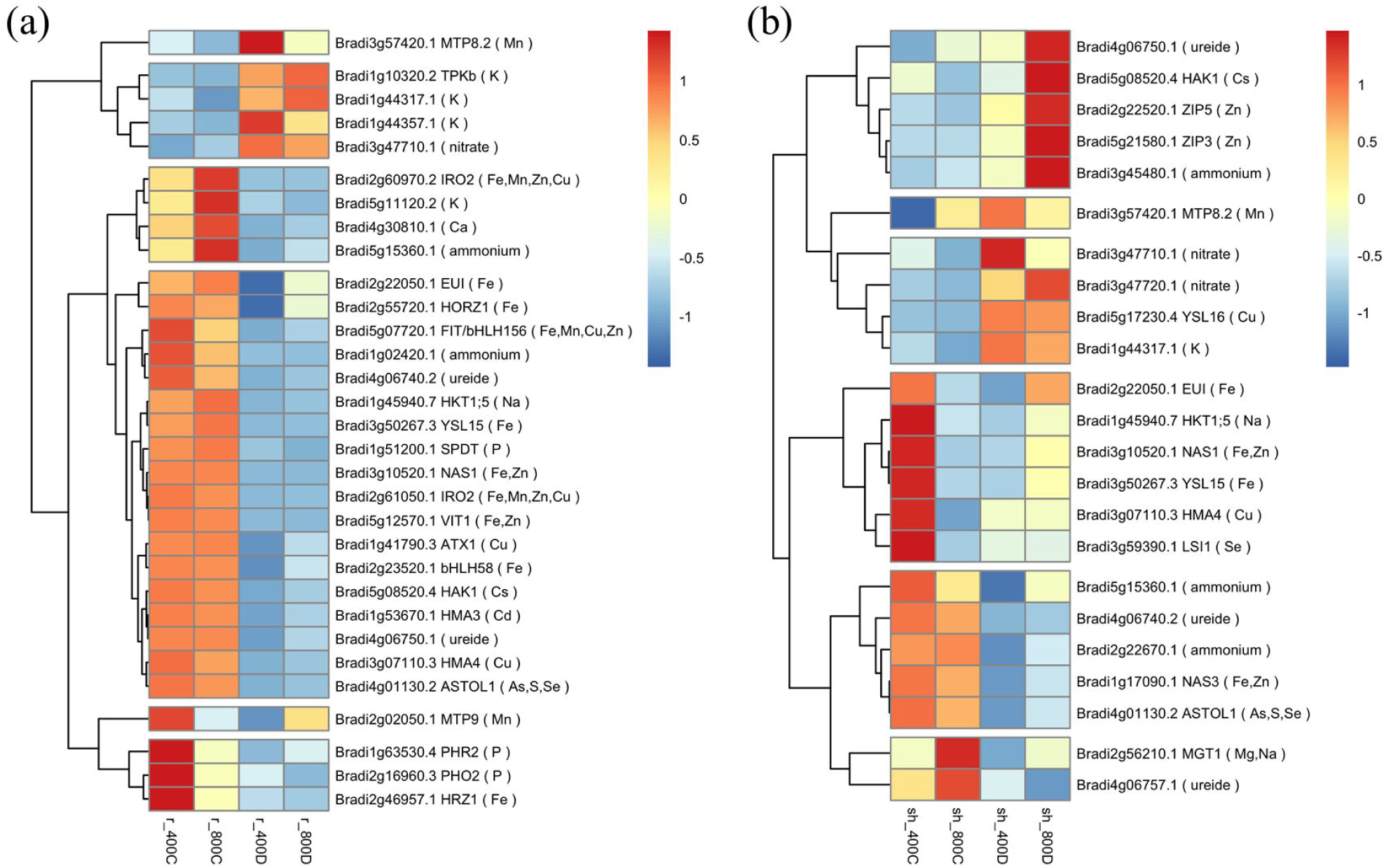
Heatmap showing the regulation of transport-related genes and known ionomic genes under control (400C), drought (400D), elevated CO_2_ (800C), and combination (800D) conditions. The log_2_ fold-change (log2(FC)) of each gene with a change greater than 1 is clustered for (a) roots and (b) shoots. The colors of the boxes represent upregulated (red) and downregulated (blue) genes. sh, shoots; r, roots.

In roots, N (ammonium)-associated genes (Figure 3a) were mostly upregulated, while the net N uptake decreased with elevated CO_2_ (Figure S5f). However, under the combined stresses of drought and elevated CO_2_ conditions, genes associated with N (nitrate), K, and Mn were generally upregulated, while the net uptake of N, K, and Mn decreased. K, P, Fe, and Mn transport-related genes were mostly downregulated under elevated CO_2_. In these cases, their net uptake was higher in elevated CO_2_ plants than in controls (Figure S7). In contrast, although the net Ca, K, Na, P, Fe, Mn, Cu, and Zn uptake decreased under drought and elevated CO_2_, their corresponding transporter genes were still found to be downregulated.

In shoots, both nitrate- and ammonium-associated genes (Figure 3b) were generally upregulated, while the partitioning of N into the shoot decreased under drought and elevated CO_2_ (Figure S5d). A distinct pattern was observed for Na transport-related genes, which were mostly downregulated, despite an increase in Na partitioning under drought and elevated CO_2_ (Figure S8d). By contrast, the partitioning of K and Fe was increased under drought and elevated CO_2_ (Figure S8), with the corresponding transporter genes found to be upregulated (Figure 3b).

### Metabolomic adjustments occur in plants’ environmental responses

Using stringent mass, RT, and fragmentation patterns, a total of 12,632 features (in positive mode) and 12,212 features (in negative mode) were observed using untargeted LC–QToF–MS. Consistent with the observed transcriptomic responses (Figure S9a), PCA revealed that PC1 explained 29.94 and 35.71% of the metabolic variation between samples in positive and negative mode, respectively, differentiating root and shoot samples. PC2 accounted for 13.48 and 9.24% (for positive and negative modes, respectively) of variation, delineating the environmental treatment conditions (water regimes, elevated CO_2_; Figure S11). Strikingly, the metabolism profile of elevated CO_2_ in roots fell into the same cluster as other root samples from drought treatments, suggesting that alterations in metabolite levels could potentially serve as an indicator of environmental stresses in the root.

439 shoot metabolites and 442 root metabolites (Figure S12), both in positive mode, were selected based on KEGG COMPOUND annotations for downstream analysis. (The *in silico* tools used herein are known to be more populated with data on the positive ionization mode (Tripathi et al., 2021)). Clustering analysis separated the metabolites into six distinct clusters reflecting relative metabolite contents across the drought, elevated CO_2_, and their interaction (Figure S12). In particular, the metabolites in shoot Cluster 3 were less abundant under drought but underwent minor changes under elevated CO_2_ compared to the control in shoots (Figure S12). Metabolites in Cluster 3 are enriched in the Citrate cycle (tricarboxylic acid cycle) and Glycolysis/Gluconeogenesis (Figure S12). Consistent with the metabolite accumulation dynamics, genes associated with photosynthesis processes show similar patterns, being down-regulated under drought (Cluster 3 in Figure S10).

### Cysteine and methionine metabolism is enhanced by elevated CO_2_ under drought

To illustrate integration of transcriptome and metabolome results, we examined the metabolism of S-containing amino acids and related compounds in plants, highlighting the regulatory function of S-adenosylmethionine (SAM). Transcript abundance of homocysteine methyltransferase (HMT; *Bradi1g13290*; EC 2.1.1.14), a key protein in the SAM cycle, was upregulated in response to drought, and more strongly so in combination with elevated CO_2_. Conversely, under drought, elevated CO_2_ decreased the expression of cystathionine γ-synthase (CGS; *Bradi1g61260*; EC 2.5.1.48), whose activity is regulated by feedback-inhibition mediated by SAM (Onouchi et al., 2004; Onouchi et al., 2005). Consistent with upregulation of SAM synthesis by drought in combination with elevated CO_2_, total SAM levels were increased by drought, and elevated CO_2_ strengthened this effect (Figure 4; S15).

**Figure 4.**
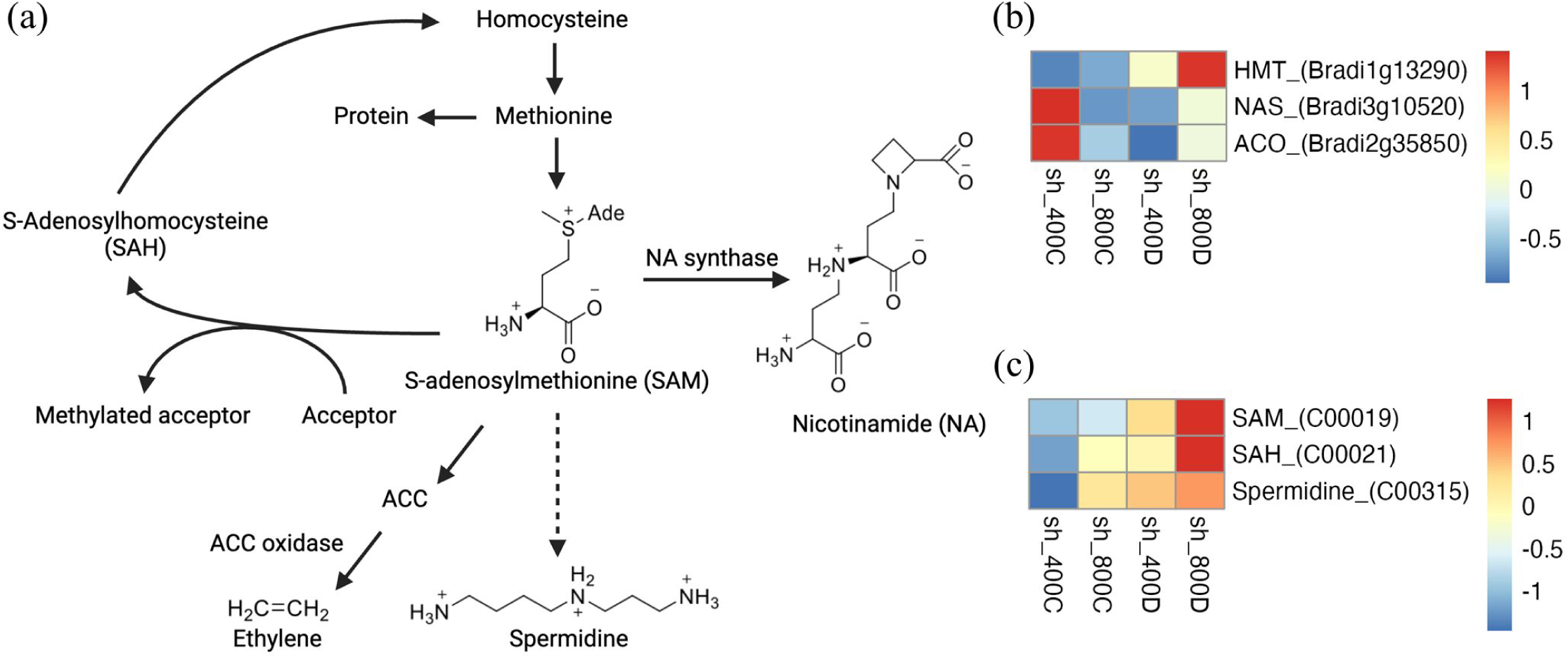
Genes and metabolites involved in the SAM cycle in plants. (a) Schematic representation of SAM metabolism. (b) The TPM values of each DEG and (c) the ion intensities (peak areas) of each metabolite were normalized using z-scores. A color gradient indicates their relative expression or accumulation levels, with red and blue representing higher and lower expression, respectively. Plants grown under control (400C), drought (400D), elevated CO_2_ (800C), or combination (800D) conditions, shoots, sh; roots, r. ACC, 1-aminocyclopropane-1-carboxylic acid; ACO, ACC oxidase; DEGs, differentially expressed genes; HMT, homocysteine S-methyltransferase; NA, nicotinamide, SAH, S-adenosylhomocysteine; SAM, S-adenosylmethionine; TPM, transcript per million.

SAM is an active form of Met and an important S-containing metabolite as a methyl donor for a range of methylation reactions; SAM is also a precursor of nicotinamide (NA), polyamine, and ethylene biosynthesis (Watanabe et al., 2021). NA, a chelator of metal cations including Fe ions, is synthesized by the trimerization of three SAM molecules, a reaction catalyzed by NA synthase (NAS; *Bradi3g10520*; EC 2.5.1.43). Drought stress downregulated NAS expression, but to a smaller extent under elevated CO_2_, which potentially stimulates the synthesis of NA from SAM. SAM is also involved in stress responses through SAM-derived polyamines(Groppa and Benavides, 2008; Alcázar et al., 2010) and ethylene. Predominant polyamines in plants, such as triamine spermidine, are synthesized by sequential additions of aminopropyl groups (derived from SAM) to putrescine via SPDS (spermidine synthase). Our findings showed that spermidine levels increased in response to drought and elevated CO_2_ compared to control conditions. Meanwhile, a gene encoding 1-aminocyclopropane-1-carboxylic acid (ACC) oxidase (ACO; *Bradi2g35850*; EC 1.14.17.4), an important enzyme in ethylene biosynthesis, was also upregulated by drought under elevated CO_2_, whereas elevated CO_2_ alone decrease its transcript abundance (Figure 4; S15).

### Metabolome annotation using CANOPUS

Spectral matching using public databases provided structural information for a small percentage of queried metabolites (3.5% in positive mode and 6.8% in negative mode). To annotate additional metabolites, we employed CANOPUS (Dührkop et al., 2021), a deep learning-based method, to predict metabolite structural classes corresponding to MSI Level 3 identifications (Sumner et al., 2007). CANOPUS classifies compounds into the multilabel and hierarchical ChemOnt ontology, i.e., ClassyFire(Djoumbou Feunang et al., 2016), which is analogous to using GO for gene sets. Since ChemOnt is multilabel, peaks may receive multiple annotations at each level; here, we report only the largest substructure of each peak. Out of the 727 and 316 fragmented peaks in positive and negative mode, respectively, 685 (94%) and 312 (98%) of the filtered features with a posterior probability greater than 0.5 (Figure S15) were annotated by CANOPUS into 16 Superclasses (File S3.1). Eight Superclasses were represented in both positive and negative mode, all within the kingdom of organic compounds (File S3.1; S3.2). Among these, lipids and lipid-like molecules have the most peaks in positive mode, while organic acids and derivatives dominate in negative modes. We primarily show results of positive mode below, but present annotations of negative mode data in the Supporting Information (Figure S16; S17).

### Annotated compound classes highlight the central role of lipids in stress-induced responses

We next used filtered CANOPUS annotations at the Class level to assess metabolites under applied stresses. To address the potential bias where large classes (e.g., 1000 metabolites) may exhibit stress-specific trends similar to smaller classes (e.g., 10 metabolites), thereby masking the relative importance of specific classes, we implemented a selection criterion. Classes were included for further analysis if their average abundance changes exceeded the 70th percentile or below the 30th percentile of all stress-induced peak area changes (Mahood et al., 2023). These selected classes were then subjected to hierarchical clustering analysis to uncover patterns of stress-induced perturbations (Figure 5). Among them, lipids and lipid-like molecules, organo-oxygen compounds, and benzenoids are the most prominent Classes.

**Figure 5.**
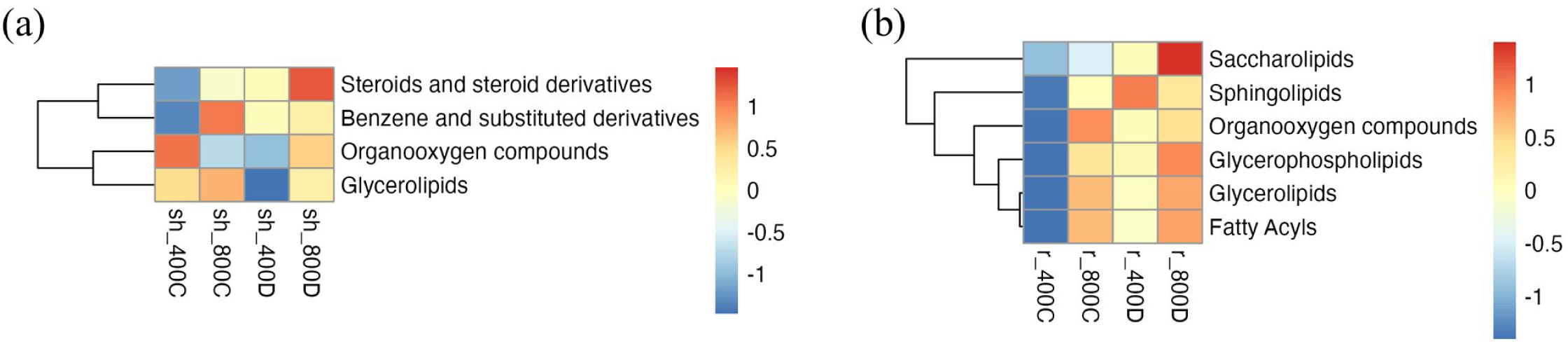
Treatment-Induced shifts of CANOPUS classes, Positive Modes. Heatmap showing the abundance changes of metabolite peaks for (a) shoots and (b) roots under control (400C), drought (400D), elevated CO_2_ (800C), and combination (800D) stresses. To be plotted, a Class’s average value must exceed the 70th percentile or fall below the 30th percentile of all treatment-induced peak area changes. The colors of the boxes represent upregulated (red) and downregulated (blue) genes. sh, shoots; r, roots.

Elevated CO_2_ induced glycerolipids (GLs) and steroids in shoots under drought (Figure 5a). In roots, we also found high perturbations in lipid classes, including glycerolipids, GPs, sphingolipids, and fatty acyls (Figure 5b). These changes may result from alterations in membrane composition, common under drought conditions (Martin et al., 1986), and/or production of lipid signaling molecules, such as sphingolipids (Berkey et al., 2012).

## Discussion

Atmospheric CO_2_ enrichment has the potential to boost C_3_ crop productivity through the so-called CO_2_ fertilization effect (Thompson et al., 2017; Tausz-Posch et al., 2020). An unexpected outcome, however, is that elevated CO_2_ often leads to lowered concentrations of key nutrients in most C_3_ plant tissues (Penuelas and Matamala, 1993; Jonard et al., 2015; Saban et al., 2019; Mariem et al., 2020; Penuelas et al., 2020). This altered nutrient status of plants may be due to a simple dilution effect caused by the increase in biomass and C content of the tissues (Poorter et al., 1997; Taub and Wang, 2008; Tausz et al., 2017). However, past studies reveal contrasting results regarding the magnitude of the growth response to elevated CO_2_, with some showing strong positive effects (McCarthy et al., 2010; Ainsworth and Long, 2021), while others report little to no response, or even negative effects, particularly under water-limited conditions (Reich et al., 2014; Gray et al., 2016).

Here, we address whether changes of nutrient concentration under combined drought and elevated CO_2_ are linked to the biomass fertilization effect of elevated CO_2_ (Hypothesis 1). We found that reduced water supply eliminates the elevated CO_2_ fertilization effect in Brachypodium, likely because interacting limitations on C, water, and nutrient assimilation under drought constrain the stimulation of plant growth. Thus, while dilution may partially explain apparent nutrient decreases at elevated CO₂, it does not fully account for the complex nutrient changes observed in shoots and roots under combined drought and elevated CO_2_.

A second hypothesis is that the negative impact of elevated CO_2_ on plant mineral status results from a reduction in transpiration-driven nutrient mass flow, caused by CO_2_-induced reductions in stomatal conductance (*g*_s_) (Del Pozo et al., 2007; Jauregui et al., 2015; Houshmandfar et al., 2018). This effect is anticipated to be greater in plants grown under drought conditions (Hypothesis 2). However, our results showed that drought and elevated CO_2_ do not equally influence the uptake and transport of all nutrients. Together with the observed reduction in growth induced by drought and elevated CO_2_ (Hypothesis 1), these results imply that drought and elevated CO_2_ alter nutrient uptake and translocation via unique processes, potentially involving multiple molecular components including transporters, channels, and chelators, (Hypothesis 3).

### Reduced water supply eliminates the elevated CO_2_ fertilization effect in Brachypodium

In response to drought, *g*_s_ reduction decreases transpiration rate and *A*_n_ (Figure 1a-c) in Brachypodium, mirroring past studies (Chaves, 1991; Flexas and Medrano, 2002; Medrano et al., 2002). Elevated CO_2_ predominantly decreases *g*_s_ and slightly stimulates *A*_n_, thereby increasing iWUE (Figure 1e). However, the stimulation of *A*_n_ is much lower than expected from first principles, likely due to acclimation responses to elevated CO_2_ over the course of development (Long et al., 2004). With extended (days to weeks) exposure to elevated CO_2_, *V*_c,max_ is often reduced through reduced Rubisco content and nitrogen investment(Drake et al., 1997; Parry et al., 2003). This response leads to decreased leaf nitrogen (Figure 2b), increased leaf C/N ratio (Figure 2c), and reduced *A*_sat_ (Figure S1f), which aligns with findings from other studies (Ainsworth and Long, 2005; Leakey et al., 2009; Ainsworth and Long, 2021).

Improved water relations under elevated CO_2_, achieved through reduced water consumption, have been reported to sustain *A*_n_ during dry periods(Ainsworth and Rogers, 2007; Soussana and Lüscher, 2007; Robredo et al., 2010; Bernacchi and VanLoocke, 2015; Deryng et al., 2016). In this study, observed *g*_s_ values were lower in the elevated CO_2_ and drought treatments, along with *V*_c,max_, *J*_max_ and *A*_sat_ downregulation which, in combination, decrease *A*_n_ (Figure 1a). According to the Farquhar et al. model (Farquhar et al., 1980), low values of *J*_max_ and, consequently of *A*_sat_, reflect reduced RuBP regeneration, while low *V*_c,max_ values are indicative of decreased Rubisco activity. The low ratio of *J*_max_ to *V*_c,max_ (Figure S1c) further implies a reduced rate of RuBP regeneration, pointing to metabolic impairment (Centritto et al., 2003). Similar findings, where elevated CO_2_ does not offset reduced carbon assimilation under drought, have been reported (Reich et al., 2014; Gray et al., 2016; Obermeier et al., 2017).

Other studies report improved water status of droughted plants under elevated CO_2_ by reducing *g*_s_, which reduces water loss and maintains high RWC (Ritchie et al., 1990; Robredo et al., 2007; Jiang et al., 2021; Wang et al., 2022). However, we found that drought stress reduced RWC regardless of CO_2_ concentration (Figure S4a); the decreased *g*_s_ under combined stresses suggests that there may be other mechanisms that negated the expected benefit on leaf water status. For example, root biomass production was reduced by elevated CO_2_ when plants were subjected to water stress (Figure S4e), possibly limiting their water acquisition capacity. These findings suggest that stomatal control and morphological adjustments jointly influence leaf water status under elevated CO_2_ and drought(Jiang et al., 2021), with elevated CO_2_ having little effect on RWC.

With CO_2_ enrichment, the C_i_/C_a_ ratio decreased (Figure 1d). This indicates acclimation of stomata towards an increased iWUE, although there was still an increase in C_i_ under CO_2_ enrichment which could increase C gain (Figure 2). Elevated CO_2_ improved iWUE to a greater extent in drought than in well-watered conditions (Figure 1e), likely due to changes in size, density, and closure response of stomata under combined stresses (Figure 1b). In parallel to decrease *g*_s_, the unchanged or decreased C_i_ and the decrease in the C_i_/C_a_ ratio observed under both drought and elevated CO_2_ treatments (Figure 1d; S2e), and resulting large drop in *A*_n_ (Figure 1a), suggest that diffusional resistance set the limit to photosynthesis rate(Cornic, 2000; Centritto et al., 2003). This implies that while elevated CO_2_ improved iWUE under drought stress, it did not ameliorate the adverse effects of drought on *A*_n_.

When water availability declined, the C/N ratio was increased in the elevated CO_2_ treatment (Figure 2), suggesting that some of the additional assimilated C was allocated to leaf growth without a corresponding allocation of N. Nitrogen limitation likely constrains the potential for increased productivity in response to elevated CO_2_ (Finzi et al., 2002; Luo et al., 2004; Hungate et al., 2006; Reich et al., 2006) but, under dry conditions, it may be difficult to separate this effect from the effect of reduced water availability.

### Reduced transpiration and changes in nutrient uptake and allocation under drought and elevated CO_2_

An alternative explanation for reduced shoot nutrient status is that elevated CO_2_ triggers stomata to close (Long et al., 2004; Xu et al., 2016), leading to lower stomatal conductance and leaf transpiration; this reduction is further accentuated by drought (Figure 1b,c). Under elevated CO_2_ alone, transpiration was reduced by 56.2%, while under the combined effects of drought and elevated CO_2_, it decreased by 93.8% (Figure 1c). Reduced transpiration may affect root nutrient uptake capacities through reduced mass flow in the soil as well as nutrient accumulation in shoots through diminished translocation via the xylem sap (Leakey et al., 2006; Houshmandfar et al., 2018). Despite these reductions, with the exception of N, the total amount of nutrients taken up by the plants in our study was increased in response to elevated CO_2_ (Figure S5; S7). However, under combined elevated CO_2_ and drought conditions, the uptake of all nutrients decreased (Figure S7). We also observed significant differences in partitioning of nutrients between shoots and roots: more N was allocated to the root at control watering and elevated CO_2_, while N allocation to the shoot was suppressed at drought and elevated CO_2_ (Figure S5d). Interestingly, despite the expected reduction in root-to-shoot nutrient translocation, there was an unexpected increase in the partitioning of Mg, K, Na, Fe, and Mn to the shoots under drought and elevated CO_2_ conditions (Figure S8). This suggests that while reduced transpiration does influence nutrient uptake and translocation, it is not the predominant factor.

While the total amount of nutrient uptake by our experimental plants is increased in response to elevated CO_2_ (Figure S7), it often does not keep pace with the stimulation of biomass production (Figure S4g). As a result, the concentration of many nutrients declined, on a dry weight basis, when grown in elevated CO_2_ (Figure 2; S7), as found previously (Wang et al., 2013; Feng et al., 2015; Tausz-Posch et al., 2020; Wang and Wang, 2021). Total nutrient uptake can decrease under elevated CO_2_, particularly when the effect of elevated CO_2_ on growth is constrained by concurrent environmental stresses (Li et al., 2016). It is possible that the reduction in transpiration-driven nutrient mass flow is offset by water limitation also restricting dry mass accumulation under elevated CO_2_, potentially balancing the reduced nutrient accumulation in the plant. Therefore, it is challenging to determine whether reductions in mineral uptake, or overall mineral content in tissues, are a direct effect of drought or simply a consequence of reduced growth.

Moreover, while transpiration is often inhibited by drought and/or elevated CO_2_, these factors do not necessarily affect root uptake and long-distance transport of nutrients via a common mechanism (Shaner and Boyer, 1976; Schulze and Bloom, 1984; Tanner and Beevers, 1990). In *Festuca*, N uptake did not correlate with changes in transpiration(Gastal and Saugier, 1989) and in maize, transpiration had no impact on ion partitioning (Tanner and Beevers, 1990). Nonetheless, recent studies identified a potential impairment of nutrient translocation to the shoot under elevated CO_2_, linking this effect to nutrient concentration in xylem sap. For example, concentrations of Ca and Mg in the xylem sap of wheat decrease under elevated CO_2_ compared to ambient CO_2_ (Houshmandfar et al., 2015), a phenomenon that cannot be solely attributed to reduced transpiration. Although the mechanisms leading to reduced concentrations in the xylem remain unclear, our findings also indicate less partitioning of Ca and Mg into the shoot under elevated CO_2_ compared to ambient CO_2_ in well-watered conditions (Figure S8).

### Shifts of gene expression and metabolite accumulation involved in transport and metal ion chelation

We found a negative impact of elevated CO_2_ on N acquisition, as both net N uptake and N tissue content were strongly decreased under applied stresses (Figure 2; S6). The reduction in N concentration at elevated CO_2_ is widely reported (Stitt and Krapp, 1999; BassiriRad et al., 2001; Tausz-Posch et al., 2020; Gojon et al., 2023), though variable(Stitt and Krapp, 1999; BassiriRad et al., 2001; Tausz-Posch et al., 2020). Further, the effect of elevated CO_2_ on plant N uptake may differ depending on the form of N used by plants (Rubio-Asensio and Bloom, 2017). Indeed, elevated CO_2_ appears to affect root nitrate uptake more negatively than it does root ammonium uptake (Jackson and Reynolds, 1996; Shimono and Bunce, 2009; Jayawardena et al., 2017; Cott et al., 2018; Ma et al., 2018; Jayawardena et al., 2021). In our study, elevated CO_2_ did not induce the expression of genes corresponding to *NRT3.1* nitrate transporter in the root, while ammonium transporter genes either showed no change (*Bradi1g02420*) or were stimulated (*Bradi5g15360*) (Figure 3). Interestingly, we observed that the relative content of spermidine, a SAM-derived polyamine, was elevated under elevated CO_2_ and/or drought conditions compared to the control (Figure 4; S18). Spermidine is a regulatory metabolite in stress responses (Watanabe et al., 2021), and exogenous spermidine application has been shown to enhance plant ammonium assimilation under drought stress(Dong et al., 2022). These results suggest a potential link between spermidine and N metabolism in plants under elevated CO_2_ and/or drought conditions.

We observed partitioning of Mg, K, Na, Fe, and Mn to the shoots under drought and elevated CO_2_ conditions (Figure S8). Accumulation of K and Na has been linked to enhanced ability to retain water (Gaxiola et al., 2001). Such an adaptive response may be reflected in the increased expression of *Bradi1g44317*, a predicted K transporter, both at ambient and elevated CO_2_. Mg is the metal center in the chlorophyll molecule and thus central to photosynthesis. Similarly, Fe and Mn are critical for the water-splitting and electron transfer reactions in photosynthesis (Suorsa and Aro, 2007). Since the photosynthetic efficiency of plants is impaired during water limitation and elevated CO_2_, accumulation of these nutrients in the shoot may reflect a strategy to maintain or improve photosynthetic efficiency (Marschner, 1995).

The case of Fe is more intriguing because we found that elevated CO_2_ strongly increased Fe partitioning in shoots under drought (Figure S8i), possibly driven by increased gene expression of *NAS1* and *YSL15* in the shoot, compared to the drought or elevated CO_2_ alone (Figure 3; 4). NAS1 catalyzes the trimerization of SAM to form one molecule of NA, which is a chelator of several transition metals, mainly Fe(II), and also Mn(II), Cu(II), and Zn(II) in all higher plants (Anderegg and Ripperger, 1989). *YSL15* transports ligands and Fe, including Fe(II)-NA(Lee et al., 2009) and it is expected that these genes are differentially regulated by Fe status of the plant. Here, plants were grown under Fe-replete conditions, and no changes in shoot Fe concentrations were observed under drought or elevated CO_2_ alone (Figure 2d). This aligns with the expectation that Fe-deficiency-induced responses, such as the expression of *NAS* and *YSL*, would not be triggered under Fe-sufficient conditions (Figure 3; 4). However, the observed increase in Fe content and gene expression under the combined conditions of drought and elevated CO_2_ indicates the involvement of other specific mechanisms during these stress conditions. A possible explanation is that elevated CO_2_ may enhance Fe chelation-based strategies in plants under drought. This enhancement could result from the increased availability of C when plants are exposed to CO_2_-enriched atmospheres (the CO_2_ fertilization effect), providing more C for synthesizing chelators. Indeed, our metabolic analysis reveals that SAM, the key precursor of NA, accumulated at a high level under drought and elevated CO_2_ (Figure 4; S15).

Moreover, we found sphingolipids, which are associated with both drought response and ionome regulation (Chao et al., 2011; Huby et al., 2020), were upregulated in the root under drought, elevated CO_2_, and the combined stresses (Figure 5). It has been shown that Arabidopsis *tsc10a* mutant reduces sphingolipid biosynthesis, which regulates ion transport through root membranes, resulting in changes in shoot elemental content(Chao et al., 2011). These findings suggest that the high accumulation of sphingolipids induced by drought and/or elevated CO_2_ may help protect against ionome imbalances.

### Conclusion

Climate change variables including precipitation (amount and distribution) and atmospheric CO_2_ concentrations are expected to alter agricultural productivity patterns worldwide and shift crop nutrient demand and usage. Existing knowledge of plant mineral nutrition and, thus, soil fertility management is inadequate to meet these challenges. Here, we provide a framework for how plant nutrient uptake and translocation processes respond to drought and elevated CO_2_ levels, integrating physiological, molecular, and non-targeted metabolic perspectives to elucidate the underlying mechanisms. We demonstrate that the elemental composition and metabolomic responses of plants to elevated CO_2_ strongly depend on water availability, and that responses differ in shoots and roots. These insights offer valuable guidance for optimizing agrochemical applications to enhance resource use efficiency and support sustainable agricultural practices under changing environmental conditions.

## Materials and Methods

### Experimental design

Seeds of *Brachypodium distachyon* Bd-21 were sterilized in 10% (v/v) household bleach for 5 min, and rinsed five times with Milli-Q water. After stratification at 4°C for 7 days, plants were grown for 7 days on 0.7% phytagel (Sigma-Aldrich, St. Louis, MO, USA) containing half-strength Murashige and Skoog (MS) medium (Murashige and Skoog, 1962) to achieve uniform seedling growth. The seedlings were then transplanted to 600ml of Profile porous ceramic rooting medium (Profile Products, Buffalo Grove, IL, USA) in Deepot D40H pots (650ml; Stuewe & Sons, Tangent, OR, USA). Plants were bottom-watered every other day using fresh water, and irrigated once weekly with water supplemented with fertilizer (1:50 dilution of Liquid Grow Plant Food; Dyna-Gro, Richmond, CA, USA).

Following 21 days of initial growth, 80 pots were randomly assigned to treatment groups comprising two-way factorial combinations of CO_2_ concentration (400 or 800 ppm) and water availability (drought or well-watered; see Supplementary Methods), resulting in 20 replicates (pots) per CO_2_ concentration × water treatment combination. At the end of the treatment period, plants were removed from their respective soils and cut into roots and shoots. All samples were flash-frozen in liquid nitrogen and then stored at −70°C.

### Measurements of Leaf Gas Exchange and Chlorophyll Fluorescence

Gas exchange and fluorescence measurements were performed on 4-week-old leaves using a LI-6800 Portable Photosynthesis System (Li-COR Biosciences Inc., Lincoln, NE, USA) equipped with a Multiphase Flash Fluorometer (6800-01A) chamber. Details of the Li-Cor settings are provided in the Supplementary Methods.

### Whole plant measurements

Half of the sampled plants were harvested for above and below-ground biomass, specific leaf area (SLA), relative water content (RWC), total carbon (C), total nitrogen (N), δ^13^C, δ^15^N, and mineral nutrient content. Details of these measurements are provided in the Supplementary Methods. The remaining plants were harvested and prepared for transcriptomic and metabolomic assays.

### RNA preparation, transcriptome sequencing, and gene expression analysis

Total RNA from root and shoot samples was extracted using the Spectrum Plant Total RNA Kit (Sigma-Aldrich, USA). cDNA libraries were prepared using the 3′ digital gene expression (3′DGE) method (Soumillon et al., 2014). Standard cDNA libraries were generated, quantified by qPCR, and sequenced on the AVITI system (Element Biosciences, San Diego, CA) at MIT BioMicro Center. All reads were mapped to the *Brachypodium distachyon* Bd21 primary transcriptome sequence (Bdistachyon_556_v3.2.transcript_primaryTranscriptOnly, downloaded from Phytozome) and the annotated sequence of the whole Bd21 plastome, which is available at Genbank (LT558597 accession) using BOWTIE2 (Langmead and Salzberg, 2012). Differential expression gene (DEG) analysis identified DEGs in response to drought, elevated CO_2_, and their interaction, using a *P*- value < 0.01 and |log_2_FC| > 1 cutoff. Heat maps were constructed using the ‘pheatmap’ package (Kolde, 2019). Gene ontology (GO) enrichment and Kyoto Encyclopedia of Genes and Genomes (KEGG) pathway enrichment of DEGs were generated using the functions ‘enrichGO’ and ‘enrichKEGG’, respectively (package: clusterProfiler (Wu et al., 2021)), with a *P*-value < 0.01 and q-value < 0.05 cutoff. DGEs related to ionome (known ionomic genes, KIGs (Whitt et al., 2020)) or those assumed to be involved in transport of a given nutrient were further analyzed. Additional details are described in the Supplementary Methods.

### Metabolite extraction and sample preparation for UHPLC-QTOF-MS analysis

Root and shoot samples were freeze-dried and ground to a fine powder using the FastPrep-24 System (MP Biomedicals, Santa Ana, CA, USA), operated at a speed of 6 m s^-1^ for 20 sec. Fresh weights of the samples were measured to obtain 50mg of dry weight for each tissue. Metabolites were extracted using a mixture of acetonitrile, isopropanol, and water (ratio of 2:2:1) containing 0.1% (v/v) formic acid. After adding the solvent mixture, samples were vortexed repeatedly over a 15-minute interval. After centrifugation for 10 min at 16,000g to remove particulates, the samples were transferred into amber HPLC vials and stored at −70°C. Liquid chromatography-mass spectrometry (LC-MS) analysis was performed using an Agilent Infinity 1260 LC system coupled to an Agilent 6545 quadrupole time-of-flight (QTOF) (Agilent, Santa Clara, CA) mass spectrometer at the Department of Chemistry Instrumentation Facility (DCIF) at MIT. Additional details are provided in the Supplementary Methods.

### Metabolomic data processing and visualization

RAW LC−MS data was converted into mzML using ProteoWizard v 3.0.7230. Total ion chromatograms (TICs) were generated for all files of a given polarity using XCMS (Mahieu et al., 2016) (Figure S18). The files for a given mode (positive or negative) were then imported into MS-DIAL v4.48(Tsugawa et al., 2015) for peak deconvolution and alignment. The LC−MS/MS spectral database from NIST17 (https://chemdata.nist.gov) and MassBank of North America (https://mona.fiehnlab.ucdavis.edu) was used for mass spectral library searches. For each polarity, MS-DIAL outputs a quantitative alignment file, displaying the peak areas of all metabolites in all samples, and a Mascot Generic Format (mgf) file of all fragmented metabolites. Detected metabolites were filtered, imputed, and normalized using R scripts (Mahood et al., 2023) in R v4.0.4 (Team, 2020). The resulting files were further processed with CANOPUS (Dührkop et al., 2021) for peak annotation, with GNPS (Nothias et al., 2020) and MS-DIAL for peak identification, as well as with MS-FINDER v3.44 (Tsugawa et al., 2016) for molecular networking. Each step is described in more detail in the Supplementary Methods. This workflow followed the pipeline developed by Elizabeth et al. (2023) (Mahood et al., 2023), with slight modifications.

### Integrative analysis of transcriptome and metabolome

Mapping of both genes and metabolites to metabolic pathways was done with the pathview package(Luo and Brouwer, 2013). Genes encoding enzymes and metabolites mediating primary amino acid metabolites were extracted from KEGG pathways for Cysteine and methionine metabolism (map00270).

### Statistical analyses

A randomized block design was implemented following a 2 x 2 factorial scheme with two atmospheric CO_2_ concentrations (400 and 800 ppm), two water regimes (well-watered and drought), and three biological replicates. The data was subjected to a two-way repeated-measures analysis of variance (ANOVA), and the means were compared using Tukey’s test at a 5% significance level using R (v4.0.4) (Team, 2020).

### Supporting Information

This section provides details on plant growth conditions, the soil drying treatment, leaf-level gas exchange and chlorophyll fluorescence measurements, whole plant measurements, RNA preparation, transcriptome sequencing, and differential gene expression analysis. It also includes the establishment of the transport-related and KIG list, untargeted metabolomic profiling of plant samples using UHPLC-QTOF-MS, pre-processing of metabolomic data, metabolic pathway enrichment analysis, peak annotation with CANOPUS, and the confirmation of CANOPUS classification. Supporting results cover plant physiology, transcriptomics, and metabolomic responses to drought and elevated CO_2_.

## Author contribution

HFC, DLD: conceptualization; HFC, TC: data collection; HFC: writing - original draft; HFC, TC, DLD: writing - review & editing; DLD: supervision; DLD: funding acquisition

## Acknowledgments

This work was supported by the MIT Climate Grand Challenges program. The authors extend their gratitude to Dr. Yilin Zhang and members of the Des Marais group for feedback.

## Reference

Ainsworth EA, Long SP (2005) What have we learned from 15 years of free-air CO2 enrichment (FACE)? A meta-analytic review of the responses of photosynthesis, canopy properties and plant production to rising CO2. New phytologist 165: 351–372

Ainsworth EA, Long SP (2021) 30 years of free-air carbon dioxide enrichment (FACE): What have we learned about future crop productivity and its potential for adaptation? Global change biology 27: 27–49

Ainsworth EA, Rogers A (2007) The response of photosynthesis and stomatal conductance to rising [CO2]: mechanisms and environmental interactions. Plant, cell & environment 30: 258–270

Alcázar R, Altabella T, Marco F, Bortolotti C, Reymond M, Koncz C, Carrasco P, Tiburcio AF (2010) Polyamines: molecules with regulatory functions in plant abiotic stress tolerance. Planta 231: 1237–1249

Anderegg G, Ripperger H (1989) Correlation between metal complex formation and biological activity of nicotianamine analogues. Journal of the Chemical Society, Chemical Communications: 647–650

BassiriRad H, Gutschick VP, Lussenhop J (2001) Root system adjustments: regulation of plant nutrient uptake and growth responses to elevated CO2. Oecologia 126: 305–320

Berkey R, Bendigeri D, Xiao S (2012) Sphingolipids and plant defense/disease: the “death” connection and beyond. Frontiers in plant science 3: 68

Bernacchi CJ, VanLoocke A (2015) Terrestrial ecosystems in a changing environment: a dominant role for water. Annual review of plant biology 66: 599–622

Catalan P, López-Álvarez D, Díaz-Pérez A, Sancho R, López-Herránz ML (2016) Phylogeny and evolution of the genus Brachypodium. Genetics and genomics of Brachypodium: 9–38

Cattivelli L, Rizza F, Badeck F-W, Mazzucotelli E, Mastrangelo AM, Francia E, Marè C, Tondelli A, Stanca AM (2008) Drought tolerance improvement in crop plants: an integrated view from breeding to genomics. Field crops research 105: 1–14

Centritto M, Loreto F, Chartzoulakis K (2003) The use of low [CO2] to estimate diffusional and non-diffusional limitations of photosynthetic capacity of salt-stressed olive saplings. Plant, Cell & Environment 26: 585–594

Champolivier L, Merrien A (1996) Effects of water stress applied at different growth stages to Brassica napus L. var. oleifera on yield, yield components and seed quality. European Journal of Agronomy 5: 153–160

Chao D-Y, Gable K, Chen M, Baxter I, Dietrich CR, Cahoon EB, Guerinot ML, Lahner B, Lü S, Markham JE (2011) Sphingolipids in the root play an important role in regulating the leaf ionome in Arabidopsis thaliana. The Plant Cell 23: 1061–1081

Chaves M (1991) Effects of water deficits on carbon assimilation. Journal of experimental Botany 42: 1–16

Cornic G (2000) Drought stress inhibits photosynthesis by decreasing stomatal aperture–not by affecting ATP synthesis. Trends in plant science 5: 187–188

Cott GM, Caplan JS, Mozdzer TJ (2018) Nitrogen uptake kinetics and saltmarsh plant responses to global change. Scientific Reports 8: 5393

Del Pozo A, Pérez P, Gutiérrez D, Alonso A, Morcuende R, Martínez-Carrasco R (2007) Gas exchange acclimation to elevated CO2 in upper-sunlit and lower-shaded canopy leaves in relation to nitrogen acquisition and partitioning in wheat grown in field chambers. Environmental and Experimental Botany 59: 371–380

Deryng D, Elliott J, Folberth C, Müller C, Pugh TA, Boote KJ, Conway D, Ruane AC, Gerten D, Jones JW (2016) Regional disparities in the beneficial effects of rising CO2 concentrations on crop water productivity. Nature Climate Change 6: 786–790

Djoumbou Feunang Y, Eisner R, Knox C, Chepelev L, Hastings J, Owen G, Fahy E, Steinbeck C, Subramanian S, Bolton E (2016) ClassyFire: automated chemical classification with a comprehensive, computable taxonomy. Journal of cheminformatics 8: 1–20

Dong L, Li L, Meng Y, Liu H, Li J, Yu Y, Qian C, Wei S, Gu W (2022) Exogenous spermidine optimizes nitrogen metabolism and improves maize yield under drought stress conditions. Agriculture 12: 1270

Drake BG, Gonzàlez-Meler MA, Long SP (1997) More efficient plants: a consequence of rising atmospheric CO2? Annual review of plant biology 48: 609–639

Dührkop K, Nothias L-F, Fleischauer M, Reher R, Ludwig M, Hoffmann MA, Petras D, Gerwick WH, Rousu J, Dorrestein PC (2021) Systematic classification of unknown metabolites using high-resolution fragmentation mass spectra. Nature biotechnology 39: 462–471

Farooq M, Wahid A, Kobayashi N, Fujita D, Basra SM (2009) Plant drought stress: effects, mechanisms and management. Sustainable agriculture: 153–188

Farquhar GD, Ehleringer JR, Hubick KT (1989) Carbon isotope discrimination and photosynthesis. Annual review of plant physiology and plant molecular biology 40: 503–537

Farquhar GD, von Caemmerer Sv, Berry JA (1980) A biochemical model of photosynthetic CO 2 assimilation in leaves of C 3 species. planta 149: 78–90

Feng Z, Rütting T, Pleijel H, Wallin G, Reich PB, Kammann CI, Newton PC, Kobayashi K, Luo Y, Uddling J (2015) Constraints to nitrogen acquisition of terrestrial plants under elevated CO 2. Global change biology 21: 3152–3168

Finzi AC, DeLucia EH, Hamilton JG, Richter DD, Schlesinger WH (2002) The nitrogen budget of a pine forest under free air CO 2 enrichment. Oecologia 132: 567–578

Flexas J, Medrano H (2002) Drought-inhibition of photosynthesis in C3 plants: stomatal and non-stomatal limitations revisited. Annals of botany 89: 183–189

Gargallo-Garriga A, Sardans J, Pérez-Trujillo M, Oravec M, Urban O, Jentsch A, Kreyling J, Beierkuhnlein C, Parella T, Peñuelas J (2015) Warming differentially influences the effects of drought on stoichiometry and metabolomics in shoots and roots. New Phytologist 207: 591–603

Gastal F, Saugier B (1989) Relationships between nitrogen uptake and carbon assimilation in whole plants of tall fescue. Plant, Cell & Environment 12: 407–418

Gaxiola RA, Li J, Undurraga S, Dang LM, Allen GJ, Alper SL, Fink GR (2001) Drought- and salt-tolerant plants result from overexpression of the AVP1 H+-pump. Proceedings of the National Academy of Sciences 98: 11444–11449

Gojon A, Cassan O, Bach L, Lejay L, Martin A (2023) The decline of plant mineral nutrition under rising CO2: physiological and molecular aspects of a bad deal. Trends in Plant Science 28: 185–198

Gray SB, Dermody O, Klein SP, Locke AM, Mcgrath JM, Paul RE, Rosenthal DM, Ruiz-Vera UM, Siebers MH, Strellner R (2016) Intensifying drought eliminates the expected benefits of elevated carbon dioxide for soybean. Nature Plants 2: 1–8

Groppa M, Benavides M (2008) Polyamines and abiotic stress: recent advances. Amino acids 34: 35–45

Hoekstra FA, Golovina EA, Buitink J (2001) Mechanisms of plant desiccation tolerance. Trends in plant science 6: 431–438

Houshmandfar A, Fitzgerald GJ, O’Leary G, Tausz-Posch S, Fletcher A, Tausz M (2018) The relationship between transpiration and nutrient uptake in wheat changes under elevated atmospheric CO2. Physiologia Plantarum 163: 516–529

Houshmandfar A, Fitzgerald GJ, Tausz M (2015) Elevated CO2 decreases both transpiration flow and concentrations of Ca and Mg in the xylem sap of wheat. Journal of plant physiology 174: 157–160

Hu Y, Burucs Z, von Tucher S, Schmidhalter U (2007) Short-term effects of drought and salinity on mineral nutrient distribution along growing leaves of maize seedlings. Environmental and Experimental Botany 60: 268–275

Huby E, Napier JA, Baillieul F, Michaelson LV, Dhondt-Cordelier S (2020) Sphingolipids: towards an integrated view of metabolism during the plant stress response. New Phytologist 225: 659–670

Hungate BA, Johnson DW, Dijkstra P, Hymus G, Stiling P, Megonigal JP, Pagel AL, Moan JL, Day F, Li J (2006) Nitrogen cycling during seven years of atmospheric CO2 enrichment in a scrub oak woodland. Ecology 87: 26–40

Jackson R, Reynolds H (1996) Nitrate and ammonium uptake for single-and mixed-species communities grown at elevated CO 2. Oecologia 105: 74–80

Jauregui I, Aroca R, Garnica M, Zamarreño ÁM, García-Mina JM, Serret MD, Parry M, Irigoyen JJ, Aranjuelo I (2015) Nitrogen assimilation and transpiration: key processes conditioning responsiveness of wheat to elevated [CO2] and temperature. Physiologia plantarum 155: 338–354

Jayawardena DM, Heckathorn SA, Bista DR, Mishra S, Boldt JK, Krause CR (2017) Elevated CO2 plus chronic warming reduce nitrogen uptake and levels or activities of nitrogen-uptake and-assimilatory proteins in tomato roots. Physiologia Plantarum 159: 354–365

Jayawardena DM, Heckathorn SA, Boldt JK (2021) A meta-analysis of the combined effects of elevated carbon dioxide and chronic warming on plant% N, protein content and N-uptake rate. AoB Plants 13: plab031

Jiang M, Kelly JW, Atwell BJ, Tissue DT, Medlyn BE (2021) Drought by CO2 interactions in trees: a test of the water savings mechanism. New Phytologist 230: 1421–1434

Jonard M, Fürst A, Verstraeten A, Thimonier A, Timmermann V, Potočić N, Waldner P, Benham S, Hansen K, Merilä P (2015) Tree mineral nutrition is deteriorating in Europe. Global change biology 21: 418–430

Kolde R (2019) Pheatmap: pretty heatmaps. R package version 1: 726

Langmead B, Salzberg SL (2012) Fast gapped-read alignment with Bowtie 2. Nature methods 9: 357–359

Leakey AD, Ainsworth EA, Bernacchi CJ, Rogers A, Long SP, Ort DR (2009) Elevated CO2 effects on plant carbon, nitrogen, and water relations: six important lessons from FACE. Journal of experimental botany 60: 2859–2876

Leakey AD, Bernacchi CJ, Ort DR, Long SP (2006) Long-term growth of soybean at elevated [CO2] does not cause acclimation of stomatal conductance under fully open-air conditions. Plant, Cell & Environment 29: 1794–1800

Lee H, Calvin K, Dasgupta D, Krinner G, Mukherji A, Thorne P, Trisos C, Romero J, Aldunce P, Barret K (2023) IPCC, 2023: Climate Change 2023: Synthesis Report, Summary for Policymakers. Contribution of Working Groups I, II and III to the Sixth Assessment Report of the Intergovernmental Panel on Climate Change [Core Writing Team, H. Lee and J. Romero (eds.)]. IPCC, Geneva, Switzerland.

Lee S, Chiecko JC, Kim SA, Walker EL, Lee Y, Guerinot ML, An G (2009) Disruption of OsYSL15 leads to iron inefficiency in rice plants. Plant physiology 150: 786–800

Lesk C, Rowhani P, Ramankutty N (2016) Influence of extreme weather disasters on global crop production. Nature 529: 84–87

Li X, Jiang D, Liu F (2016) Soil warming enhances the hidden shift of elemental stoichiometry by elevated CO2 in wheat. Scientific Reports 6: 23313

Li Y, Ye W, Wang M, Yan X (2009) Climate change and drought: a risk assessment of crop-yield impacts. Climate research 39: 31–46

Long SP, Ainsworth EA, Rogers A, Ort DR (2004) Rising atmospheric carbon dioxide: plants FACE the future. Annu. Rev. Plant Biol. 55: 591–628

Luo W, Brouwer C (2013) Pathview: an R/Bioconductor package for pathway-based data integration and visualization. Bioinformatics 29: 1830–1831

Luo Y, Su B, Currie WS, Dukes JS, Finzi A, Hartwig U, Hungate B, McMurtrie RE, Oren R, Parton WJ (2004) Progressive nitrogen limitation of ecosystem responses to rising atmospheric carbon dioxide. Bioscience 54: 731–739

Ma Q, Wang J, Sun Y, Yang X, Ma J, Li T, Wu L (2018) Elevated CO2 levels enhance the uptake and metabolism of organic nitrogen. Physiologia plantarum 162: 467–478

Mahieu NG, Genenbacher JL, Patti GJ (2016) A roadmap for the XCMS family of software solutions in metabolomics. Current opinion in chemical biology 30: 87–93

Mahood EH, Bennett AA, Komatsu K, Kruse LH, Lau V, Rahmati Ishka M, Jiang Y, Bravo A, Louie K, Bowen BP (2023) Information theory and machine learning illuminate large-scale metabolomic responses of Brachypodium distachyon to environmental change. The Plant Journal 114: 463–481

Mariem SB, Gámez AL, Larraya L, Fuertes-Mendizabal T, Cañameras N, Araus JL, McGrath SP, Hawkesford MJ, Murua CG, Gaudeul M (2020) Assessing the evolution of wheat grain traits during the last 166 years using archived samples. Scientific Reports 10: 21828

Marschner H (1995) Mineral nutrition of higher plants 2nd edn. Institute of Plant Nutrition University of Hohenheim: Germany

Martin BA, Schoper JB, Rinne RW (1986) Changes in soybean (Glycine max [L.] Merr.) glycerolipids in response to water stress. Plant physiology 81: 798–801

McCarthy HR, Oren R, Johnsen KH, Gallet-Budynek A, Pritchard SG, Cook CW, LaDeau SL, Jackson RB, Finzi AC (2010) Re-assessment of plant carbon dynamics at the Duke free-air CO2 enrichment site: interactions of atmospheric [CO2] with nitrogen and water availability over stand development. New Phytologist 185: 514–528

Mcgrath JM, Lobell DB (2013) Reduction of transpiration and altered nutrient allocation contribute to nutrient decline of crops grown in elevated CO2 concentrations. Plant, Cell & Environment 36: 697–705

Medrano H, Escalona JM, Bota J, Gulías J, Flexas J (2002) Regulation of photosynthesis of C3 plants in response to progressive drought: stomatal conductance as a reference parameter. Annals of botany 89: 895–905

Murashige T, Skoog F (1962) A revised medium for rapid growth and bio assays with tobacco tissue cultures. Physiologia plantarum 15

Myers SS, Zanobetti A, Kloog I, Huybers P, Leakey AD, Bloom AJ, Carlisle E, Dietterich LH, Fitzgerald G, Hasegawa T (2014) Increasing CO2 threatens human nutrition. Nature 510: 139–142

Nemani RR, Keeling CD, Hashimoto H, Jolly WM, Piper SC, Tucker CJ, Myneni RB, Running SW (2003) Climate-driven increases in global terrestrial net primary production from 1982 to 1999. science 300: 1560–1563

Nothias L-F, Petras D, Schmid R, Dührkop K, Rainer J, Sarvepalli A, Protsyuk I, Ernst M, Tsugawa H, Fleischauer M (2020) Feature-based molecular networking in the GNPS analysis environment. Nature methods 17: 905–908

Obermeier WA, Lehnert LW, Kammann C, Müller C, Grünhage L, Luterbacher J, Erbs M, Moser G, Seibert R, Yuan N (2017) Reduced CO2 fertilization effect in temperate C3 grasslands under more extreme weather conditions. Nature Climate Change 7: 137–141

Onouchi H, Lambein I, Sakurai R, Suzuki A, Chiba Y, Naito S (2004) Autoregulation of the gene for cystathionine γ-synthase in Arabidopsis: post-transcriptional regulation induced by S-adenosylmethionine. *In*. Portland Press Ltd.

Onouchi H, Nagami Y, Haraguchi Y, Nakamoto M, Nishimura Y, Sakurai R, Nagao N, Kawasaki D, Kadokura Y, Naito S (2005) Nascent peptide-mediated translation elongation arrest coupled with mRNA degradation in the CGS1 gene of Arabidopsis. Genes & development 19: 1799–1810

Parry M, Andralojc P, Mitchell RA, Madgwick P, Keys A (2003) Manipulation of Rubisco: the amount, activity, function and regulation. Journal of experimental botany 54: 1321–1333

Penuelas J, Fernández-Martínez M, Vallicrosa H, Maspons J, Zuccarini P, Carnicer J, Sanders TG, Krüger I, Obersteiner M, Janssens IA (2020) Increasing atmospheric CO2 concentrations correlate with declining nutritional status of European forests. Communications Biology 3: 125

Penuelas J, Matamala R (1993) Variations in the mineral composition of herbarium plant species collected during the last three centuries. Journal of Experimental Botany 44: 1523–1525

Peterson P, Sheaffer C, Hall M (1992) Drought effects on perennial forage legume yield and quality. Agronomy Journal 84: 774–779

Poorter H, Van Berkel Y, Baxter R, Den Hertog J, Dijkstra P, Gifford R, Griffin K, Roumet C, Roy J, Wong S (1997) The effect of elevated CO2 on the chemical composition and construction costs of leaves of 27 C3 species. Plant, Cell & Environment 20: 472–482

Reich PB, Hobbie SE, Lee T, Ellsworth DS, West JB, Tilman D, Knops JM, Naeem S, Trost J (2006) Nitrogen limitation constrains sustainability of ecosystem response to CO2. Nature 440: 922–925

Reich PB, Hobbie SE, Lee TD (2014) Plant growth enhancement by elevated CO2 eliminated by joint water and nitrogen limitation. Nature Geoscience 7: 920–924

Ritchie SW, Nguyen HT, Holaday AS (1990) Leaf water content and gas-exchange parameters of two wheat genotypes differing in drought resistance. Crop science 30: 105–111

Rivas-Ubach A, Sardans J, Pérez-Trujillo M, Estiarte M, Peñuelas J (2012) Strong relationship between elemental stoichiometry and metabolome in plants. Proceedings of the National Academy of Sciences 109: 4181–4186

Robredo A, Pérez-López U, de la Maza HS, González-Moro B, Lacuesta M, Mena-Petite A, Muñoz-Rueda A (2007) Elevated CO2 alleviates the impact of drought on barley improving water status by lowering stomatal conductance and delaying its effects on photosynthesis. Environmental and experimental botany 59: 252–263

Robredo A, Pérez-López U, Lacuesta M, Mena-Petite A, Muñoz-Rueda A (2010) Influence of water stress on photosynthetic characteristics in barley plants under ambient and elevated CO2 concentrations. Biologia Plantarum 54: 285–292

Rubio-Asensio JS, Bloom AJ (2017) Inorganic nitrogen form: a major player in wheat and Arabidopsis responses to elevated CO2. Journal of Experimental Botany 68: 2611–2625

Saban JM, Chapman MA, Taylor G (2019) FACE facts hold for multiple generations; Evidence from natural CO2 springs. Global Change Biology 25: 1–11

Schulze E-D, Bloom AJ (1984) Relationship between mineral nitrogen influx and transpiration in radish and tomato. Plant physiology 76: 827–828

Shaner DL, Boyer JS (1976) Nitrate reductase activity in maize (Zea mays L.) leaves: II. regulation by nitrate flux at low leaf water potential. Plant Physiology 58: 505–509

Shimono H, Bunce JA (2009) Acclimation of nitrogen uptake capacity of rice to elevated atmospheric CO2 concentration. Annals of Botany 103: 87–94

Singh S, Singh G, Singh P, Singh N (2008) Effect of water stress at different stages of grain development on the characteristics of starch and protein of different wheat varieties. Food Chemistry 108: 130–139

Soumillon M, Cacchiarelli D, Semrau S, van Oudenaarden A, Mikkelsen TS (2014) Characterization of directed differentiation by high-throughput single-cell RNA-Seq. BioRxiv: 003236

Soussana JF, Lüscher A (2007) Temperate grasslands and global atmospheric change: a review. Grass and forage science 62: 127–134

Stitt M, Krapp A (1999) The interaction between elevated carbon dioxide and nitrogen nutrition: the physiological and molecular background. Plant, Cell & Environment 22: 583–621

Sumner LW, Amberg A, Barrett D, Beale MH, Beger R, Daykin CA, Fan TW-M, Fiehn O, Goodacre R, Griffin JL (2007) Proposed minimum reporting standards for chemical analysis: chemical analysis working group (CAWG) metabolomics standards initiative (MSI). Metabolomics 3: 211–221

Suorsa M, Aro E-M (2007) Expression, assembly and auxiliary functions of photosystem II oxygen-evolving proteins in higher plants. Photosynthesis Research 93: 89–100

Tanner W, Beevers H (1990) Does transpiration have an essential function in long-distance ion transport in plants? Plant, Cell & Environment 13: 745–750

Taub DR, Wang X (2008) Why are nitrogen concentrations in plant tissues lower under elevated CO2? A critical examination of the hypotheses. Journal of Integrative Plant Biology 50: 1365–1374

Tausz M, Bilela S, Bahrami H, Armstrong R, Fitzgerald G, O’Leary G, Simon J, Tausz-Posch S, Rennenberg H (2017) Nitrogen nutrition and aspects of root growth and function of two wheat cultivars under elevated [CO2]. Environmental and Experimental Botany 140: 1–7

Tausz-Posch S, Tausz M, Bourgault M (2020) Elevated [CO 2] effects on crops: Advances in understanding acclimation, nitrogen dynamics and interactions with drought and other organisms. Plant Biology 22: 38–51

Team RC (2020) RA language and environment for statistical computing, R Foundation for Statistical. Computing

Thompson M, Gamage D, Hirotsu N, Martin A, Seneweera S (2017) Effects of elevated carbon dioxide on photosynthesis and carbon partitioning: a perspective on root sugar sensing and hormonal crosstalk. Frontiers in Physiology 8: 578

Tripathi A, Vázquez-Baeza Y, Gauglitz JM, Wang M, Dührkop K, Nothias-Esposito M, Acharya DD, Ernst M, van der Hooft JJ, Zhu Q (2021) Chemically informed analyses of metabolomics mass spectrometry data with Qemistree. Nature chemical biology 17: 146–151

Tsugawa H, Cajka T, Kind T, Ma Y, Higgins B, Ikeda K, Kanazawa M, VanderGheynst J, Fiehn O, Arita M (2015) MS-DIAL: data-independent MS/MS deconvolution for comprehensive metabolome analysis. Nature methods 12: 523–526

Tsugawa H, Kind T, Nakabayashi R, Yukihira D, Tanaka W, Cajka T, Saito K, Fiehn O, Arita M (2016) Hydrogen rearrangement rules: computational MS/MS fragmentation and structure elucidation using MS-FINDER software. Analytical chemistry 88: 7946–7958

Vandegeer R, Miller RE, Bain M, Gleadow RM, Cavagnaro TR (2012) Drought adversely affects tuber development and nutritional quality of the staple crop cassava (Manihot esculenta Crantz). Functional Plant Biology 40: 195–200

Wang L, Feng Z, Schjoerring JK (2013) Effects of elevated atmospheric CO2 on physiology and yield of wheat (Triticum aestivum L.): a meta-analytic test of current hypotheses. Agriculture, Ecosystems & Environment 178: 57–63

Wang M, Zheng Q, Shen Q, Guo S (2013) The critical role of potassium in plant stress response. International journal of molecular sciences 14: 7370–7390

Wang Z, Wang C (2021) Magnitude and mechanisms of nitrogen-mediated responses of tree biomass production to elevated CO2: A global synthesis. Journal of Ecology 109: 4038–4055

Wang Z, Wang C, Liu S (2022) Elevated CO2 alleviates adverse effects of drought on plant water relations and photosynthesis: a global meta-analysis. Journal of Ecology 110: 2836–2849

Watanabe M, Chiba Y, Hirai MY (2021) Metabolism and regulatory functions of O-acetylserine, S-adenosylmethionine, homocysteine, and serine in plant development and environmental responses. Frontiers in Plant Science 12: 643403

Whitt L, Ricachenevsky FK, Ziegler GZ, Clemens S, Walker E, Maathuis FJ, Kear P, Baxter I (2020) A curated list of genes that affect the plant ionome. Plant Direct 4: e00272

Wu D, Shen Q, Cai S, Chen Z-H, Dai F, Zhang G (2013) Ionomic responses and correlations between elements and metabolites under salt stress in wild and cultivated barley. Plant and Cell Physiology 54: 1976–1988

Wu T, Hu E, Xu S, Chen M, Guo P, Dai Z, Feng T, Zhou L, Tang W, Zhan L (2021) clusterProfiler 4.0: A universal enrichment tool for interpreting omics data. The innovation 2

Xu Z, Jiang Y, Jia B, Zhou G (2016) Elevated-CO2 response of stomata and its dependence on environmental factors. Frontiers in plant science 7: 657

